# A novel system for maintaining *Varroa destructor* mites on artificial diets and its application for studying mites as a vector for honey bee viruses

**DOI:** 10.1101/2019.12.12.874107

**Authors:** Francisco Posada-Florez, Eugene V. Ryabov, Matthew C. Heerman, Yanping Chen, Jay D. Evans, Steven C. Cook, Daniel E. Sonenshine

**Affiliations:** USDA, ARS Bee Research Laboratory, Beltsville Agricultural Research Center, Beltsville, MD 20705, USA; Department of Biological Sciences, Old Dominion University, Norfolk, VA 23529, USA

**Keywords:** *Apis mellifera*, Iflavirus, *Varroa destructor* virus-1 (VDV1), Deformed wing virus, infectious viral cDNA clone for VDV1, DWV-B, virus vector, parafilm sachet, artificial diet, *in vitro* rearing

## Abstract

The mite *Varroa destructor* is one of the most destructive parasites of the honey bee (*Apis mellifera*) and the primary cause of colony collapse in most regions of the world. These mites cause serious injury to their hosts, especially during the larval and pupal stages, and serve as the vector for several viruses, which affect honey bee health causing colony death. Attempts by beekeepers to control these mites have yielded limited success. The inability to rear populations of mites *in vitro* that excludes contact with their honey bee hosts has stymied research of *Varroa* biology. Previous attempts to rear and/or maintain *Varroa* mites *in vitro* by feeding them on artificial diets have had limited success. Several methods were plagued by mechanical failures including leaking membranes and, thus far, none have been widely adopted. Here we report a robust system for maintaining *Varroa* mites that includes an artificial diet, which does not contain honey bee tissue-derived components, thus making it particularly valuable in studying mite vectoring of honey bee viruses. With our system we demonstrated for the first time that *Varroa* mites maintained on an artificial diet supplemented with the particles of honey bee viruses, cDNA clone-derived genetically tagged Varroa destructor virus-1 and wild-type Deformed wing virus, can acquire and later transmit these viruses to recipient honey bee pupae. Along with providing an opportunity to study parasites and pathogens in the absence of honey bee hosts, this i*n vitro* system for *Varroa* mite maintenance is both scalable and consistent. These features can be used to better understand mite nutritional needs, metabolic activity, responses to chemicals and other biological functions.

## Introduction

The ectoparasitic mite, *Varroa destructor* (Anderson and *Trueman*) (Acari: *Varroidae*), is believed to be the major factor in the widespread collapse of honey bee colonies in North America as well as many parts of Europe and Asia [1]. Attempts by beekeepers to control *Varroa* mites have had only limited success [2, 3]. Consequently, there is a need to find new methods to control these parasites without harming the host bees. Unfortunately, efforts to gain a better understanding of the biology of *Varroa* have been stymied by the lack of a ready supply of mites along with a repeatable system for maintaining and/or rearing populations of mites for research purposes. Attempts to rear *Varroa* mites on liquid artificial diets have been the subject of intense investigation since the 1980’s [4, 5]. Honey bee host pupae, the only life stage suitable for mite reproduction, are not available during the fall and winter in temperate areas, including most of the USA and Europe, and natural populations of *Varroa* mites do not arrive at sufficient levels needed for scientific studies until late summer [6]. Successful rearing, or even long-term maintenance, of mite populations would greatly facilitate research on disease transmission, metabolism, reproduction, and the evaluation of control methods.

There have been several attempts to rear populations of *Varroa* mites *in vitro* using artificial diets. Perhaps the first successful attempt to feed *Varroa* on an artificial diet was reported by Bruce et al. [4], who stretched a thin (~10 µm) parafilm membrane containing a dietary medium over modified queen cells, which are used for queen-rearing. Mites were observed to survive for up to 120 h, and many mites also laid eggs. However, this system was plagued by evaporation and contamination of the diet. Further modifications to this system using queen rearing cells covered with parafilm minimized leakage and were successful for maintaining a number of the much smaller tracheal mite, *Acarapis woodi* [7]. Later, Talbart et al. [5] produced synthetic chitosan membranes for holding an artificial diet with which to rear *Varroa* mites but this system also had limited success and has not yet been adopted by other researchers. This issue is likely due to earlier systems that required relatively complex components; only a few devices could be manufactured at a time, with the result that tests were done with very few mites [5, 8]. Attempts to reproduce these published methods revealed many setbacks (authors’ unpubl. data), especially related to membrane leakage, which drowned the mites. These physical limitations made it difficult, if not impossible to carry out repeatable tests for research. Because a system for rearing populations of *Varroa* mites *in vitro* remains elusive, a system is still needed for maintaining sufficient numbers of mites for research purposes.

The success of maintaining *Varroa* mites on artificial diets for research purposes depends upon three major issues: 1) availability of numerous healthy adult female mites; 2) the composition of the artificial diet; and 3) a device and membrane system for housing mites and containing the diet, respectively. Unfortunately, sufficient numbers of *Varroa* mites for research remain seasonally restricted. Nonetheless, for further progress in maintaining large numbers of mites *in vitro* with artificial diets, the first task is to create a simple but reliable system that can allow mites housed in a device to have access to a diet that does not leak, evaporate or is prone to contamination [9]. The second task is to provide a membrane thin enough (10 to 15 µm), to allow *Varroa* mites to reach the diet with their mouthparts, which are very short [10]. Finally, methods must contain a sufficient volume of liquid to feed mites for many days or even weeks without compromising the quality and/or integrity of the diet. In this report, we describe a robust system including a membrane based on that of Avila and colleagues [11] for maintaining large numbers of *Varroa* mites numbers on completely host-free diets. Further, we validate the system’s utility in an experiment of *Varroa* mite vectoring of honey bee viruses, specifically, the novel cDNA clone-derived *Varroa destructor virus-1* (VDV1 or DWV-B).

## Materials and Methods

### V-BRL Diet

The artificial diet, hereafter termed the V-BRL (Varroa-Bee Research Lab) diet was based on Tabart et al. and Bruce et al. [4, 5]. Briefly, the diet included 30% Schneider’s medium, 30% CMRL-1000, 1% Hanks salt solution (without NaCO_3,_ CaCl or MgSO_4_), 10% bovine serum, 4% Insect medium supplement (cell culture type), 10% TC-100 Insect medium (with glutamine and NaCO_3_) and 15% sterile water (all from Sigma-Aldrich, St. Louis MO). To minimize contamination, fresh diet was prepared in a fume hood and stored frozen (−20°C) in small aliquots. No antibiotics or antifungal agents were included to avoid compromising microbiota that may provide essential nutrients for mite nutrition, since species of *Diplorickettsia, Arsenophonus, Morganella, Spiroplasma, Enterococcus*, and *Pseudomonas* have been reported to inhabit *Varroa* mites [12].

### *Varroa* mites

The sugar roll method [13] was used to collect phoretic mites from honey bee colonies that had not received miticidal applications. Following collection, and after rinsing mites with tap water, they were placed on petri dishes lined with dry tissue paper. Mites were checked for activity prior to experimental selection, and only active mites used.

### Device and membrane

Snap-cap polypropylene 1.5 mL microcentrifuge tubes were chosen for constructing the device housing the mites (Fig 1). The tubes were cut crosswise at the point that allowed a 1 cm diameter opening. Cut surfaces were smoothed with sandpaper to avoid damaging the parafilm (American National Can Company, NY) membranes that would later be attached. Prior to using the devices, while wearing sterile nitrile gloves, containers were transferred to a laminar flow hood, disinfected for 2 min with 1.0 % bleach, followed directly by 2 min in 70 % ethanol, then 2 min rinsing in sterile water and then exposed for 1 h to ultraviolet light. Following disinfection and while still in the laminar flow hood, each device was covered with a thin sheet of sterilized parafilm stretched to 16.6±5.86 µm and wrapped over the open end of the device. Membrane thickness was checked with a Marathon digital micrometer (Marathon Corp., Canada). Diet solution (10 µL) was pipetted onto the center of the parafilm membrane, after which the membrane plus diet solution was covered with a second layer of sterile parafilm, forming the parafilm sachet [11, 14, 15]. The snap-cap lid on the other side of the device was opened to insert a female *Varroa* mite. Before closing, the lid was punctured with a sterile no. 1 stainless steel pin to form a minute hole (approximately 0.5 mm) to allow air to escape so the parafilm sachet would not rupture when the lid was snapped shut (Fig 1). The completed devices were placed in a dark incubator (Thermofisher Scientific, San Jose, CA) set at 32.1 ± 0.3 °C, and having 82.2 ± 1.3 % relative humidity.

**Figure 1.**
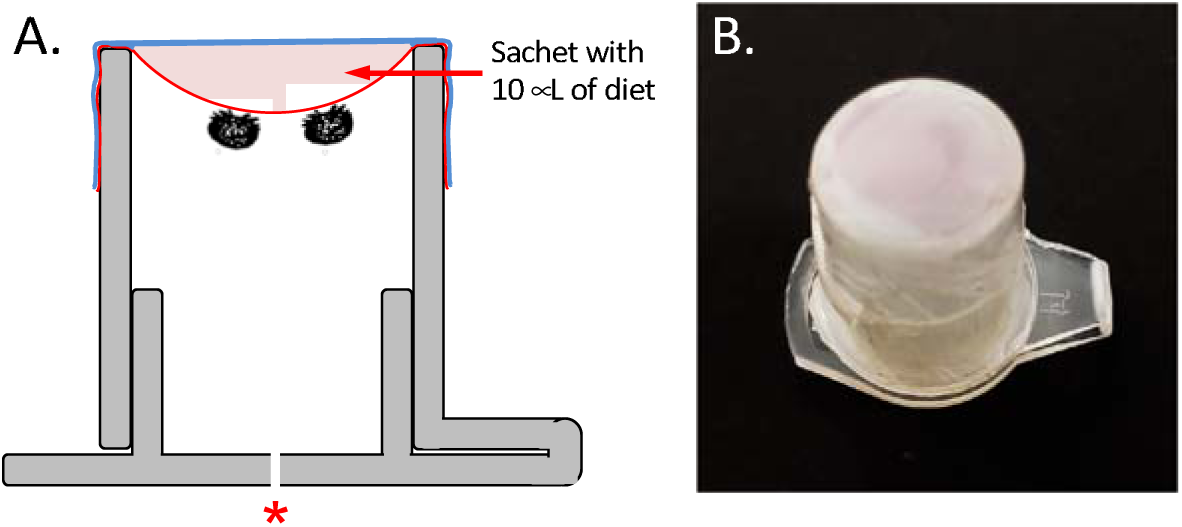
Details of the device and membrane sachet comprising the system used for maintaining *Varroa* mites during experiments. (A). Diagram showing the design of the device housing the mites. Arrow shows parafilm sachet with diet. Red asterisk indicates hole in the lid for air pressure relief. Two mites are shown attached to inner membrane to feed on diet. (B) Photo of the complete system, comprising the device housing the mites and attached parafilm sachet. The sachet is filled with 10 µL of diet solution.

### Confirming mite feeding on the artificial diet

Because mites defecate frequently when allowed to feed [16], the number of mite excretory deposits made on the parafilm membrane was recorded daily as a measure of mite feeding. Samples of diet solution contained 10^4^ sterile fluorescent isothiocyanate (FITC) labeled beads from Thermofisher (Carlsbad, CA), to determine whether the mites had fed. Samples of mites were collected after they died, decontaminated with (70% ethanol) and examined with a Zeiss Axio Imager.M2 (Dublin, CA) microscope at 488 excitation/520 emission for fluorescent beads in their tissues as described previously [17]. In addition, the number of mite excretory deposits made each day was recorded using a Zeiss Axioskop 2 Plus compound microscope (Dublin, CA) to inspect the walls of the device and membrane.

### Tests of device performance and *Varroa* mite longevity

Three trials to evaluate the system for maintaining mites were conducted. In the first trial, 10 replicates were prepared with three *Varroa* mites in each device. In the second trial, 10 replicates were prepared; six with three mites and four with 4 mites, for a total of 34 mites. In the third trial, 15 replicates were prepared; four devices with two mites, four with three mites and 7 with four mites each for a total of 42 mites. In each trial 10µL of diet plus beads were placed within the parafilm sachet. As a positive control for survival, female *Varroa* mites were allowed to feed on honey bee pupae. Pupae were collected from bee hives, placed in clear gelatin capsules (Capsuline, Pompano Beach, FL) with several mites, and mite and bee survival were monitored daily. Pupae were replaced after 3–4 days with fresh bee pupae to ensure that the pupal hosts had not deteriorated, transformed into an adult bee, or were otherwise unsuitable, as described previously [18]. As a negative control for mite survival, mites were housed in devices but membrane sachets contained only water. Replicate devices and controls were incubated as described above. Mite mortality was recorded daily. Also, devices were checked daily for leakage of the diet, or whether air bubbles or other obstructions blocked access to the diet. Finally, the diet was inspected for contamination and evaporation as described previously [9].

### Design of infectious cDNA clone of *Varroa destructor* virus-1 (VDV1) and production of clone-derived inocula

We designed a full-length infectious cDNA clone of a Californian isolate of VDV1, GenBank Accession number MN249174 (S1 Text, S2 Text). The cDNA had an introduced genetic marker, an *Asi*SI restriction site at the position 277 nt, which distinguished clone-derived VDV1 from wild-type VDV1 strains allowing us to trace transmission of this VDV1 isolate (S2 Text). The VDV1 cDNA clone and clone-derived infectious virus particles were produced using previously described approach [19], and detailed in S1. In brief, the full-length cDNA clone of the virus was produced using total RNA from honey bees sourced from California in 2016 (isolate CA-07-2016) which showed high VDV1 and low DWV levels [20]). Two overlapping cDNA fragments amplified by RT-PCR using specific primers, and the synthetic gene corresponding to the 277 nt 5’ part of the genomic RNA were assembled in a plasmid vector (S1 Table; S1 Text). The resulting full-length VDV1 cDNA plasmid construct was used to prepare the template for in *vitro* transcription. To produce clone-derived VDV1 inoculum, the purified *in vitro* RNA full-length VDV1 transcript generated using the linearized VDV1 cDNA plasmid was injected into the hemolymph of purple eye honey bee pupae. Pupae were incubated 4 days at +33°C (82.2 ± 1.3% relative humidity) to allow propagation of the clone-derived virus infection. Tissue extracts containing the clone-derived VDV1 virus particles were collected by homogenizing infected pupae in PBS, and then filtering the supernatant through a 0.22μm nylon syringe filter (ThermoFisher, Waltham, MA) to remove host-derived cellular materials. The VDV1 extract introduced to the *Varroa* diet contained the clone-derived VDV1 (1.7 × 10^8^ per µL) and background wild-type DWV derived from the recipient pupae (1.6 × 10^5^ per µL), see S1 text for additional details.

### Acquisition of viruses by *Varroa* mites from artificial diet

The filtered tissue extract introduced to the *Varroa* diet contained virus particles of the clone-derived VDV1 at a concentration of 1.7 × 10^8^ genome equivalents/μL and background wild-type DWV derived from the recipient pupae used for recovery of a clone-derived VDV1 at a concentration of 1.6 × 10^5^ genome equivalents/μL. We supplemented the artificial diet with filtered PBS tissue extract containing particles of the cDNA clone-derived VDV1 and wild-type DWV (80% diet, 20% filtered PBS tissue extract), Fig 2. The virus-supplemented diet contained 3.3×10^7^ genome equivalents of the clone-derived VDV1 and 3.2 × 10^4^ genome equivalents of wild-type DWV per µL diet. The diet supplemented with PBS (80% diet, 20% PBS) containing no tissue extract was used as a control.

**Figure 2.**
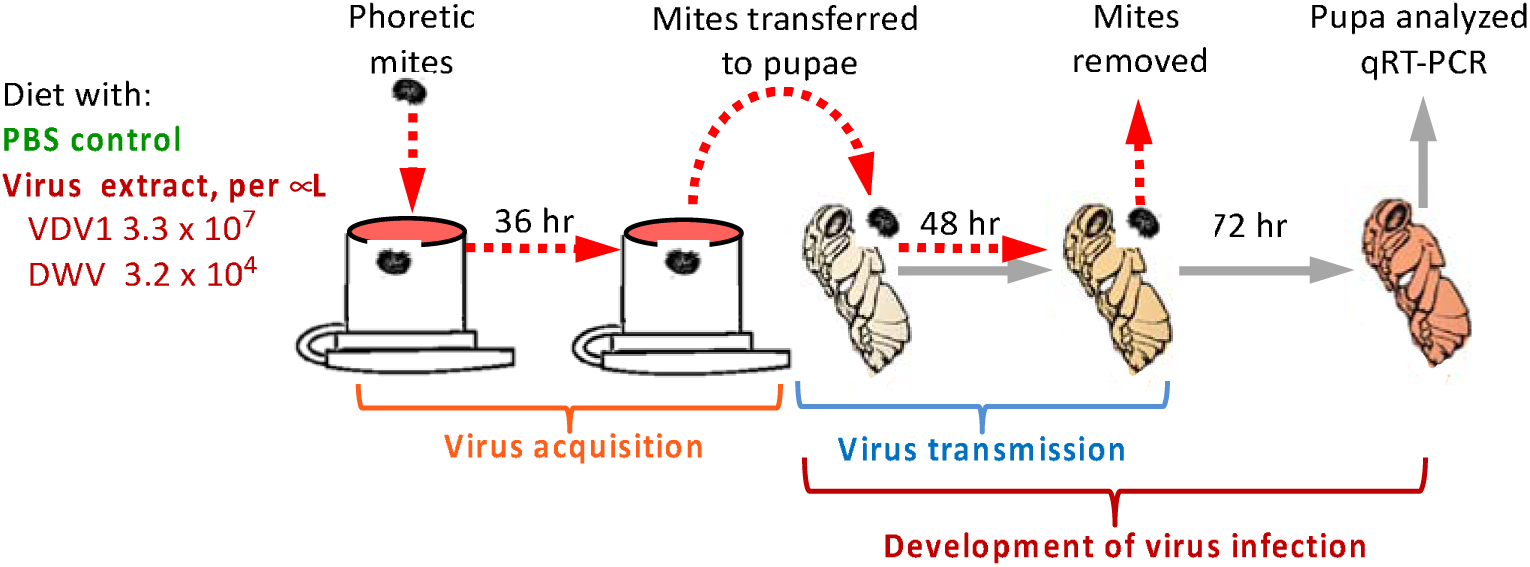
□Application of the developed system for studying the vectoring of honey bee viruses by *Varroa* mites. Schematic representation of the experimental design. Mites were allowed to feed for 30 h either on the diet contained cDNA clone derived particles of VDV1, 3.3 × 10^7□^genome equivalents per□μ□, and wild-type DWV, 3.2 × 10^4□^genome equivalents per□μL, or on the control diet containing PBS free of viral particles.

### Transmission of the acquired viruses to honey bee pupae

Following the 36h acquisition period the surviving mites were placed individually on a pink-eyed honey bee pupa housed gelatin capsules and placed in a dark incubator (see above) for 48h to allow for the transmission of the virus to the pupae. Pupae were then incubated for an additional 72h to allow virus replication (Fig 2). To determine whether viral transmission occurred, total RNA was extracted from each of pupae from the experimental (n=15) and control (n=9) treatment groups, then extracts subject to RT-qPCR to quantify copy numbers of VDV1 and DWV genomic RNA and assess transcription levels of honey bee actin by RT-qPCR using the primers described in S1 Table as in [20]. To further demonstrate that VDV1 detected in the pupae derived from the clonal cDNA virus acquired by the mites from the artificial diet, the RT-PCR fragment corresponding to the 5’ terminal 1200 nt was digested with the *Asi*SI restriction enzyme. The digested and undigested PCR fragments were separated by electrophoresis in 1.2 % agarose gel and visualized using ethidium bromide staining.

### Data analysis

The integrity of the device and diet were monitored and recorded using Zeiss Axio camera attached to a stereoscopic microscope. The survival of the mites on the diet was analyzed using a Kaplan-Meier test (JMP, version 12, SAS, Cary, NC). Other variables included the number of fecal pellets deposited per day, and the number of viral replicates in pupae after *Varroa* mite transmission were analyzed using ANOVA or *t*-tests.

## Results

### Device, membrane and mite survival

The feeding device (Fig 1) allowed *Varroa* mites to feed on the artificial diet without honey bee-derived components through a ~10 µm parafilm membrane. No evidence of diet leakage, evaporation or contamination, or other mechanical failures precluding mite’s access to the diet was recorded. We compared survival of mites from the three trials having devices with parafilm sachets containing the V-BRL artificial diet against survival of mites from both the positive and negative control groups (Fig 3; S2 Table). In contrast to diet fed-mites, most mites survived up to 25 days while feeding on bee pupae. Mite survival declined precipitously after day 10. The average survival time while feeding on pupae was 9.42 days (Fig 3, green dotted line). We found that mites began dying within 6h after confinement in the artificial feeding device without diet. Of the 32 mites held in the container without food (negative control group), all but two were dead within one day; the remaining two died by day two (Fig 3, black dotted line). Notably, mites incubated in the devices containing water deposited only minute amounts of excreta, indicating a lack of feeding. Contrary to poor survival by the mites in devices containing water (mean survival time ± standard error: 1.06 ± 0.04 days), mites survived for significantly long periods when kept in the devices containing diet (Fig 3, solid lines).

**Figure 3.**
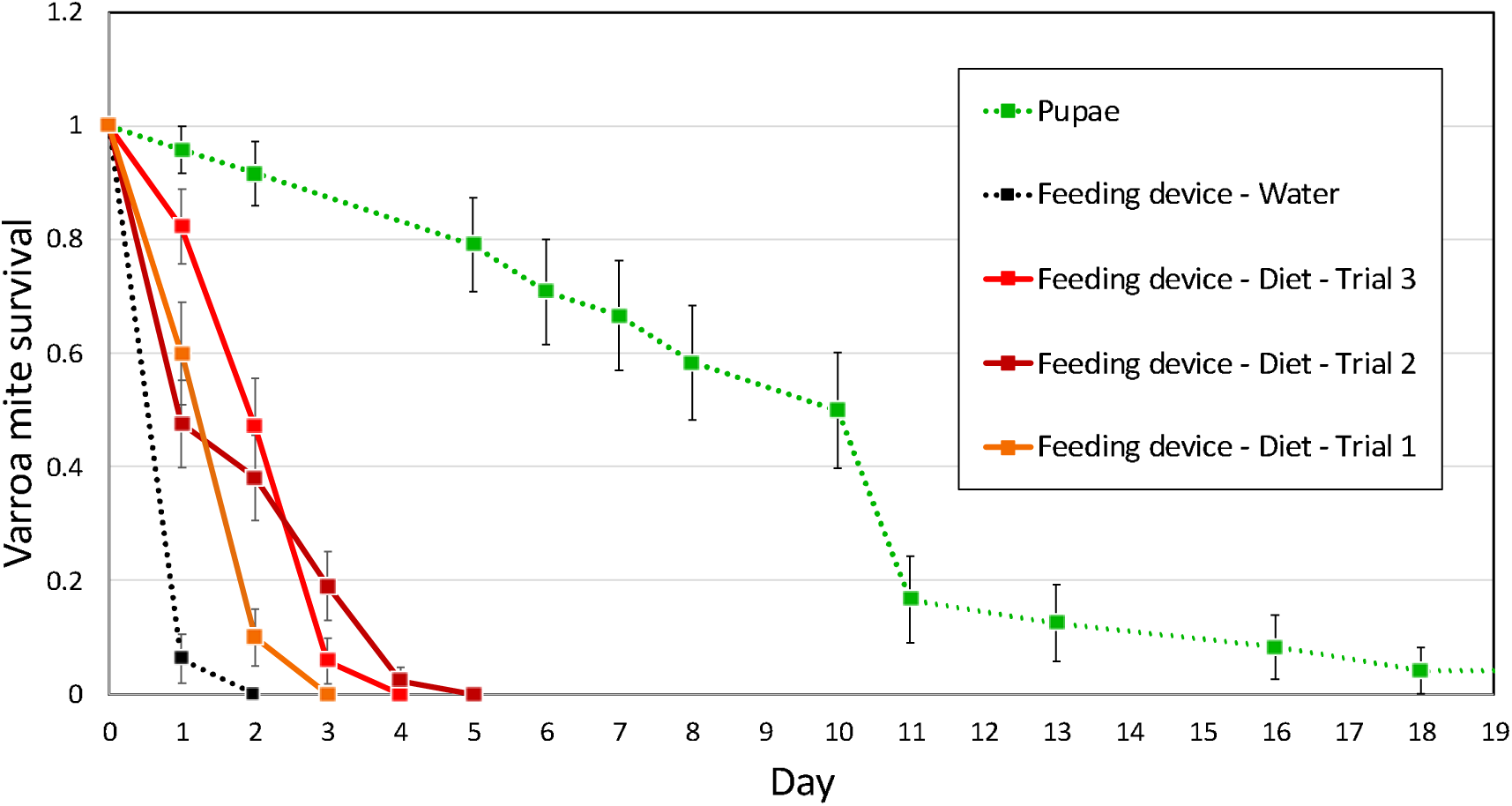
Survival of *Varroa* mites on the artificial diet during 3 separate trials compared to mites feeding on honey bee pupae and a negative control. Survival curves are shown for 3 separate diet trials (orange, red and dark red lines) compared to mites feeding on honey bee pupae (green dotted line) and mites without a food source (black dotted line). Data points on the graph represent the mean percent survival (± std. error) in days for each trial.

Three trials each involving 10 diet chambers, involving 30 to 42 mites in total in each trial, were carried out (Fig 3, Table 1; S2 Table), with mean survival times ± standard error for three trials were 1.70 ± 0.12, 2.35 ± 0.35, and 2.07 ± 0.20 days. In the first trial, mites survived for up to 4 days (d); slightly more than 60% survived 1 d, 10% survived 2 d, and the remainder survived 3 d. In the second trial, 82% survived 1 d, 48 % survived 2 d, 5% survived 3 d and 1% 4 d. In the third trial, 48% survived 1 d, 39% survived 2 d, 20% survived 3 d, 3% survived 4 d, and none survived 5 d. The longevity of mites did not depend on the number of mites housed in the devices (data not shown).

**Table 1.** Mean percent survival (± std. error) for *Varroa* mites reared on honey bee pupae, artificial diet, or water in the feeding devices. The statistical significance of the results was determined by the Wilcoxon Chi-square test (P < 0.001, Chi square = 84.49).

| **Treatment group** | **Number of mites** | **Mean survival time**<br>**(days)** | **Standard error** |
| --- | --- | --- | --- |
| Pupae (gelatin capsule) | 23 | 9.42 | 0.90 |
| Feeding device -<br>Water | 32 | 1.06 | 0.04 |
| Feeding device -<br>Diet -Trial 1 | 30 | 1.70 | 0.12 |
| Feeding device -<br>Diet -Trial 2 | 34 | 2.35 | 0.15 |
| Feeding device -<br>Diet -Trial 3 | 42 | 2.07 | 0.20 |

Evidence for mite feeding was supported by finding numerous mite excretory deposits (Fig 4A, S3 Table) and the finding of FITC-labeled fluorescent beads in excreta of the mites fed on the bead-containing diet (Fig 4B) but not in excreta for those feeding diet without fluorescent beads (Fig 4 C). The daily average deposition of excreta was 2.55 ± 0.52 deposits per mite. Statistical analysis of the excreta data for each of the four days showed that the differences in excreta per live mite per day were not statistically significant (one-way ANOVA, P = 0.3797, Fig 4 D).

**Figure 4.**
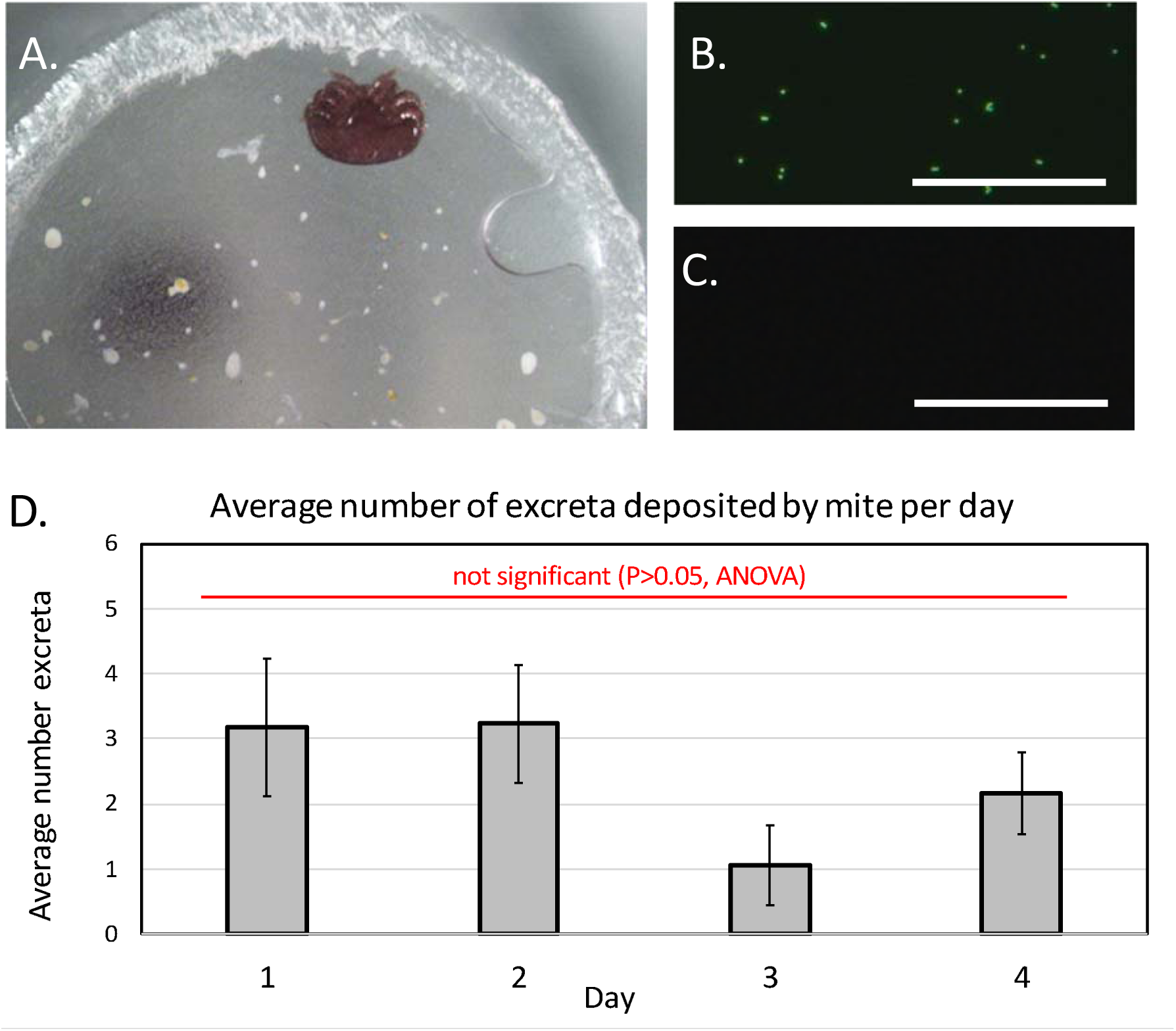
Evidential support of *Varroa* mites feeding on artificial diets. (A) Photograph shows the parafilm membrane with a live mite attached and numerous excretory deposits. (B) Micrograph showing fluorescent microbeads in mite feces collected from devices housing mites provided diet containing microbeads. (C) Micrograph showing no fluorescent microbeads in mite feces collected from devices housing mites provided diet containing no microbeads. (D) The mean (± se) total daily number of excretions deposited by mites on the membrane sachet. Measurement bar = 50 µM.

### Evidence of acquisition of viruses by mites and transmission to honey bee pupae

There was no statistically significant difference in the survival of mites fed on the diet containing the viruses versus mites fed on the control diet over a 72-h study period (S1 Fig, S4 Table, S5 Table). While there was no significant difference between the levels of honey bee actin mRNA in the treatment and control groups (P=0.1395, S6 Table), the levels of DWV and VDV1 were statistically significantly higher in pupae exposed to virus-fed mites compared to those in the control group (P<0.01 and P<0.001 respectively; Fig 5A). One-way ANOVA of the virus accumulation in the control and treatment groups were highly significant, (P<0.01) for DWV (F=11.83, df=23 P=0.00234; Cohen’s d=1.473107, large effect size.), (P<0.001) for VDV1 (F=17.95, df=23. P=0.00034, Cohen’s d=2.009988, large effect size). Importantly, high levels of DWV and VDV1 exceeding 10^9^ virus copies per pupa, were observed only in the treatment group. High levels of VDV1 were observed in 7 of 15 pupae of the treatment group pupae, while only one pupa developed high DWV levels in the control (Fig 5A). This result could be explained by a 1000-fold difference between the levels of DWV and VDV1 in the diets containing 3.2 × 10^4^ and 3.3 × 10^7^ genome equivalent per µL respectively.

**Figure 5.**
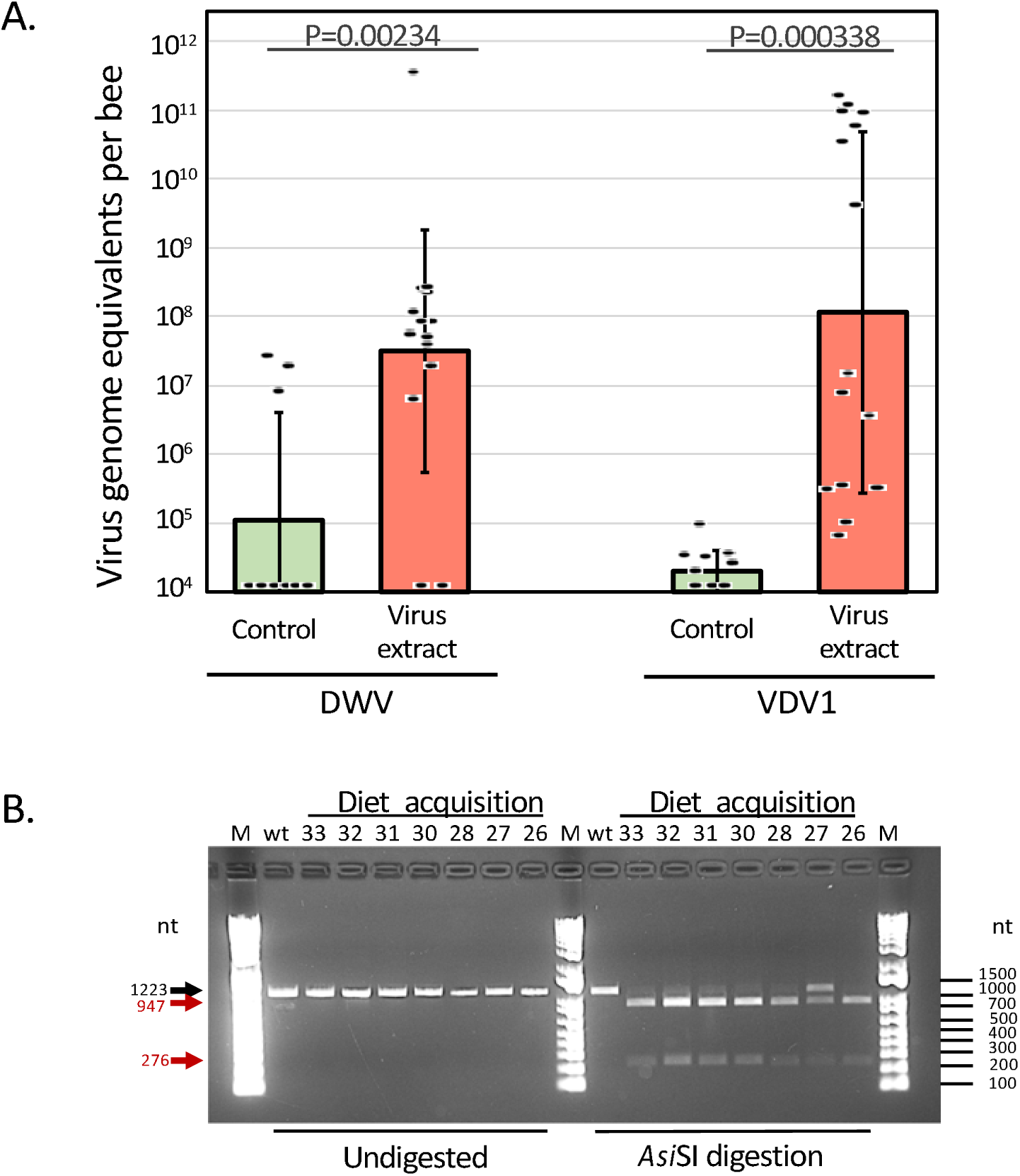
Transmission of VDV1 and DWV acquired by *Varroa* mites from artificial diet to honey bee pupa. (A) Levels of clone-derived VDV1 and wild-type DWV in honey bee pupae exposed to the *Varroa* mites which acquired virus from the diet. Black dots indicate virus load in individual honey bees. The columns show the average virus genome equivalent per µL numbers for each treatment ± SD. Statistically significant differences (unpaired Student’s t-test) are indicated above the bars. Numerical values underlying the summary graphs are provided in S6 Table. (B) To confirm the VDV1 clonal identity, the 1250 nt RT-PCR fragments corresponding to the 5’-terminal region of DWV and VDV1 RNA genomes were amplified with the primers specific to both DWV and VDV1 using RNA extracts from the experimental pupae with the virus levels exceeding 10^9^ genome copies from the “Virus extract” group (S6 Table), and from the wild-type VDV1 and DWV-infected pupae, “wt”. The undigested 1250 nt fragments (left) and *Asi*SI-digested (right). Expected fragment sizes, undigested (black arrow) and *Asi*SI-digested (red arrows), are shown on the left, DNA ladder sizes are shown on the right.

We confirmed that VDV1 derived from the artificial diet replicated in the treatment group pupae to high levels based on the diagnostic *Asi*SI restriction site (Fig 5B). This restriction site was not present in the wild-type isolates of VDV1 or DWV (Fig 5B, lane “wt”). Further evidence showing that the VDV1 transmitted to pupae was the cDNA clone VDV1 acquired from the artificial diet comes from digesting the RT-PCR fragment corresponding to the 5’ terminal 1200 nt with *Asi*SI restriction enzyme. All seven of the recipient pupae that developed high virus levels showed that the *Asi*SI site was present in VDV1 (Fig 5B). Notably, the sample isolated from the 27 pupae of the treatment group contained high levels of both DWV and VDV1 (Table S5), but only VDV1 was digested as agarose gel analysis showed (Fig 5B, lane 27).

## Discussion

Here we describe a simple, but robust, system including an artificial diet, which was free of honey bee-derived components, that successfully maintained *Varroa* mites for a study of their vectoring of honey bee viruses to host pupae. In contrast to the previous artificial systems for rearing/maintaining *Varroa* mites, we used the accumulation of mite excreta and the presence of the fluorescent microbeads acquired from the diet in the excreta to confirm that mites had successfully fed through the membrane on the artificial diet. With this system, many mites survived consistently for 4 days (rarely, 7 days Posada et al unpublished), and an average longevity for up to five days, which is similar to the longevity of mites reported using other artificial systems [5]. The finding of live mites and excretory pellets on the parafilm membranes strongly suggests that the mites were feeding repeatedly on the diet, similar to the numerous excretory pellets found by Egekwu et al [18] for mites surviving on honey bee pupae. Notably, the greatest number of excretory deposits were found in the devices in which the mites survived the longest as supported by finding FITC-labeled fluorescent beads in the mite excreta. These findings show that *Varroa* mites can not only survive on the artificial diet, but also can remain vigorous and suitable for experiments of viral transmission such as the current study showing virus acquisition and transmission from diet to pupae. Thus, the important question was not whether mites survived for long periods like those fed exclusively on pupae [18], but whether enough mites fed and survived during the period of the experiments in which they were used.

Our results with using the novel cDNA clone of genetically-tagged VDV1 also show that the acquisition period of 36 h was sufficient for *Varroa* mites feeding on the artificial diet to acquire sufficient numbers of viral particles to transmit them to the honey bee pupae, wherein they proliferated to high levels. No viral transmission to host pupae occurred subsequent to mites feeding on the control diet supplemented only with fluorescent beads. Although the results of our study showing viral transmission to honey bees by *Varroa* mites is not novel, here we show for the first time that virus particles of clonal VDV1 and DWV remained infectious in the artificial V-BRL diet long enough to be acquired by *Varroa* mites and subsequently transmitted to honey bee host pupae.

A significant challenge in this experiment was potential presence of viruses in both the recipient honey bee pupae and the *Varroa* mites sourced from the honey bee colonies maintained in apiaries known to harbor viruses. Therefore, it was also necessary to distinguish between the virus introduced through feeding on the diet and those that may have already been present either in the mites or in the recipient honey bee pupae. This challenge was resolved by using cDNA clone-derived VDV1 (also known as DWV-B) that was tagged with a rare *Asi*SI restriction enzyme site to differentiate it from the wild type (Fig 5B). The absence of pupae with high DWV and VDV1 levels (above 10^9^ genome equivalents) in the control group (Fig 5A), i.e. by mites exposed to 36 h feeding on virus-free diet, was in agreement with the recent report by which indicated that DWV is vectored by *Varroa* mites via a non-persistent mechanical route [17].

The virus acquisition and subsequent transmission experiment presented in this report provides a “proof of concept” that validates the utility of the *in vitro* system for maintaining *Varroa* mites for research purposes. We demonstrated that mites could acquire, and vector virus particles obtained from the filtered dietary suspension rather than only from living infected cells. Demonstrating mite transmission of viruses acquired from an artificial diet is particularly important in the light of the recent finding that *Varroa* mites ingest honey bee cells (including fat body cells) rather than just hemolymph [17, 22]. The described diet and experimental approach could be used to investigate *Varroa* vectoring of other honey bee viruses [23]. Feeding mites on a medium that is not derived from honey bees will provide definitive insights into which mite-vectored viruses are capable of and/or persisting in their mite carriers. Finally, an important point of the system is the relative ease and manufacture of our artificial feeding device as it provides a convenient system to test mite survival when presented with a large variety of diet formulations. This is a significant improvement on the feeding devices described previously [4, 21].

## Acknowledgments

This research was supported in part by the United States Department of Agriculture (USDA) National Institute of Food and Agriculture (NIFA) grant 2017-06481 (EVR, YPC and JDE), and ORISE fellowship (FPF) and an APHIS-ARS interagency agreement (19-8130-0745-IA) (SCC and JDE). The funders had no role in study design, data collection and analysis, decision to publish, or preparation of the manuscript. USDA is an equal opportunity provider and employer.

## Supporting information

**S1 Figure.** Mite survival analysis of the virus acquisition and transmission experiment.

**S1 Table.** Primers and the synthetic gene used in this study.

**S2 Table.** Mite survival.

**S3 Table.** Mite excreta data.

**S4 Table.** Virus acquisition and transmission experiment: Mite survival summary.

**S5 Table.** Virus acquisition and transmission experiment: Mite survival analysis.

**S6 Table.** Virus acquisition and transmission experiment, RT-qPCR quantification of VDV1, DWV and honey bee actin.

**S1 Text.** Design of the infectious cDNA clone of Varroa destructor virus-1 (VDV1) and production of the clone-derived VDV1 inoculum.

**S2 Text.** Nucleotide sequence of the Varroa destructor virus-1 infectious cDNA construct (GenBank accession number MN249174).

**S1 Video.** Varroa feeding on artificial diet artificial supplemented with the particles of honey bee viruses.

## Supporting information

**S1 Figure.**
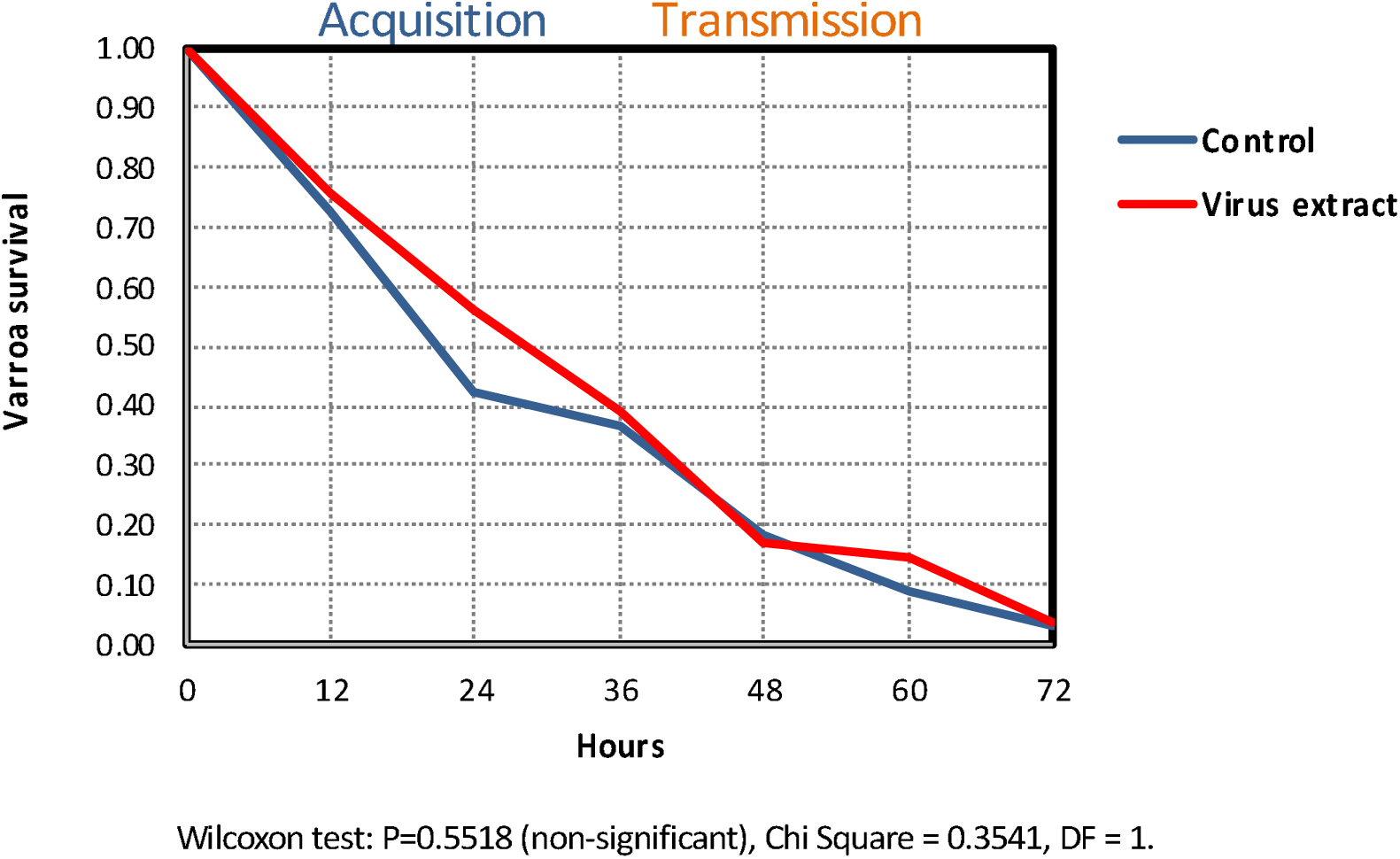
Mite survival analysis of the virus acquisition and transmission experiment.

**S1 Table.**
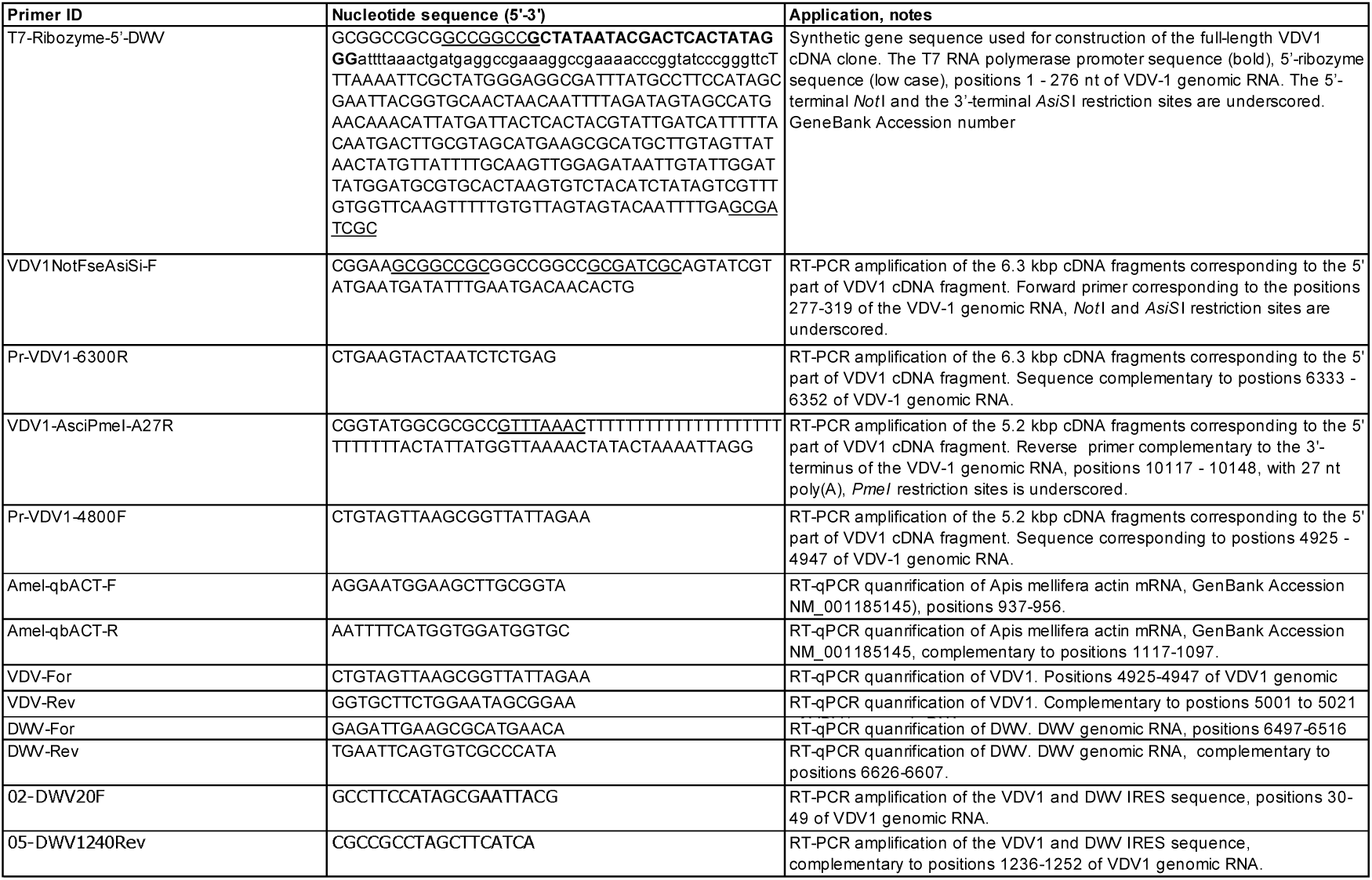
Primers and the synthetic gene used in this study (VDV1 genomic RNA - GenBank Accession MN249174, DWV genomic RNA - GenBank Accession number MG831200)

**S2 Table.**
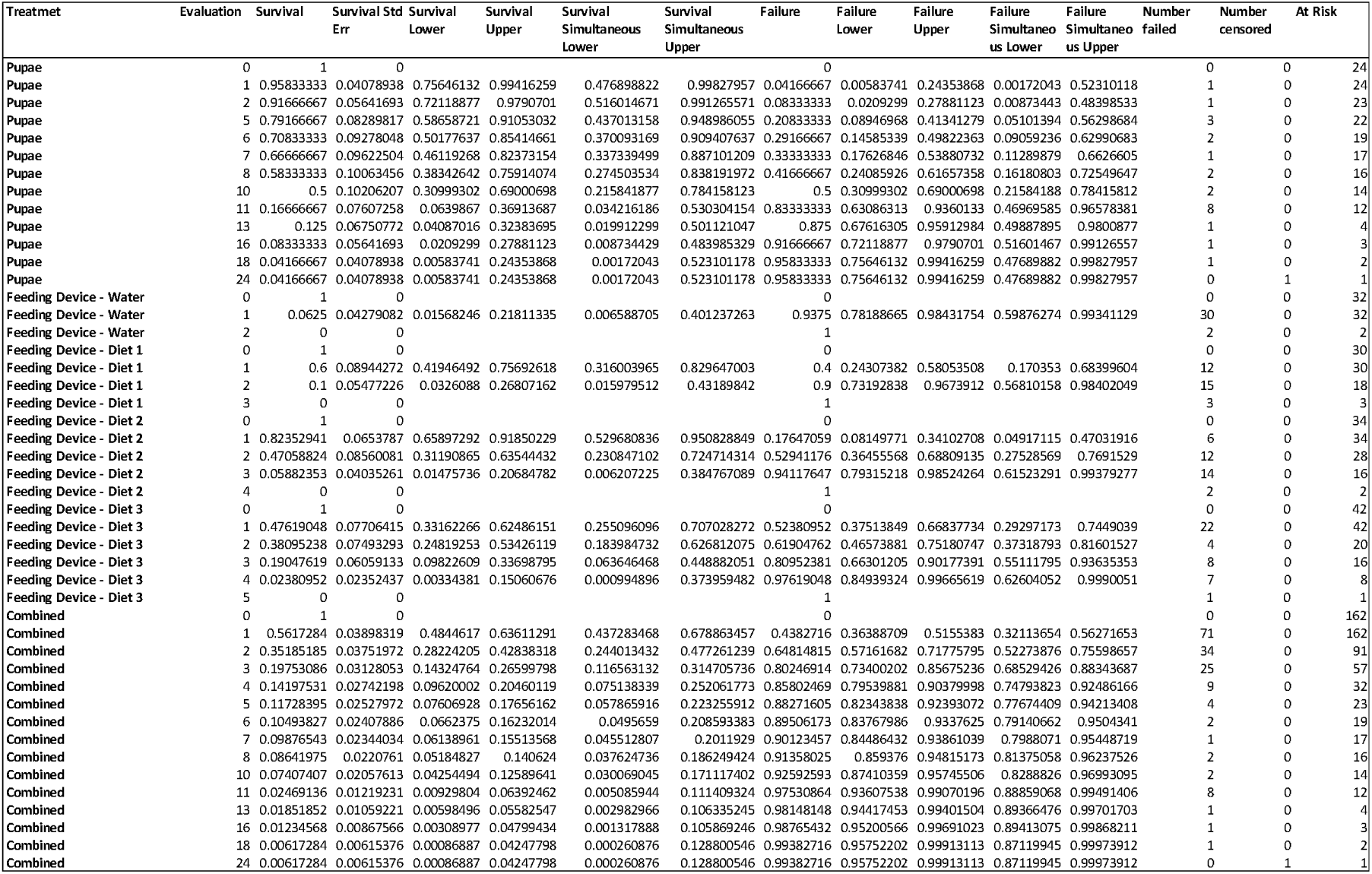
Mite survival.

**S3 Table.**
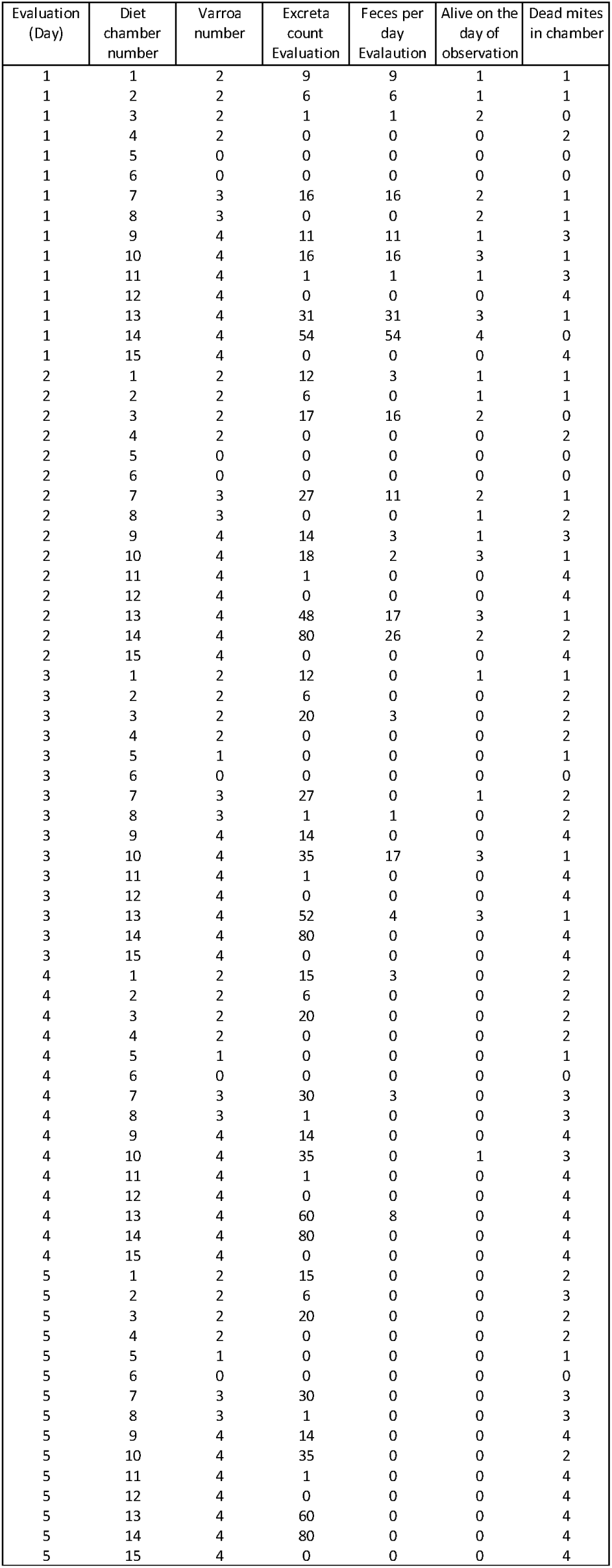
Mite excreta data.

**S4 Table.**
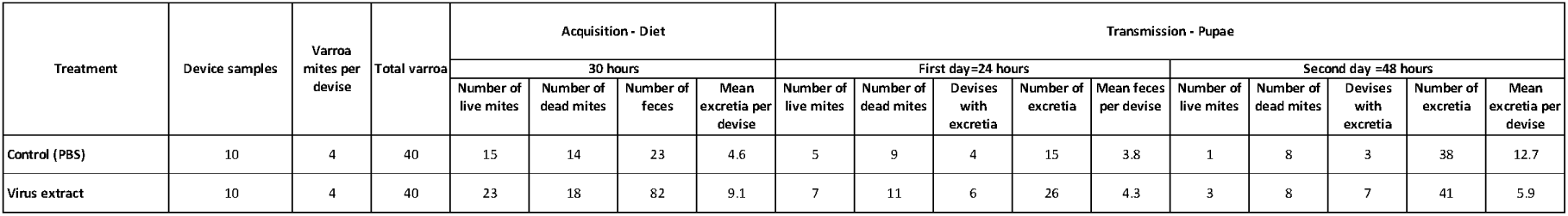
Virus acquisition and transmission experiment: Mite survival summary.

**S5 Table.** Virus acquisition and transmission experiment: Mite survival analysis.

| Treatment | Hours | Survival | SurvStdErr |
| --- | --- | --- | --- |
| Control | 0 | 1.00 | 0.00 |
| Control | 12 | 0.73 | 0.08 |
| Control | 24 | 0.42 | 0.09 |
| Control | 36 | 0.36 | 0.08 |
| Control | 48 | 0.18 | 0.07 |
| Control | 60 | 0.09 | 0.05 |
| Control | 72 | 0.03 | 0.03 |
| Virus extract | 0 | 1.00 | 0.00 |
| Virus extract | 12 | 0.76 | 0.07 |
| Virus extract | 24 | 0.56 | 0.08 |
| Virus extract | 36 | 0.39 | 0.08 |
| Virus extract | 48 | 0.17 | 0.06 |
| Virus extract | 60 | 0.15 | 0.06 |
| Virus extract | 72 | 0.04 | 0.03 |

| Summary |  |  |  |  |
| --- | --- | --- | --- | --- |
| Group | Number failed | Number censored | Mean | Std Error |
| Control | 32 | 1 | 33.4545 | 3.4955 |
| Virus extract | 38 | 3 | 36.2927 | 3.18047 |
| Combined | 70 | 4 | 35.027 | 2.3415 |

| Quantiles |  |  |  |  |  |
| --- | --- | --- | --- | --- | --- |
| Group | Median Time | Lower 95% | Upper 95% | 25% Failures | 75% Failures |
| Control | 24 | 24 | 48 | 12 | 48 |
| Virus extract | 36 | 24 | 48 | 24 | 48 |
| Combined | 30 | 24 | 36 | 12 | 48 |

| Tests between groups |  |  |  |
| --- | --- | --- | --- |
| Test | ChiSquare | DF | Prob>ChiSq |
| Log-Rank | 0.3071 | 1 | 0.5795 |
| Wilcoxon | 0.3541 | 1 | 0.5518 |

**S6 Table.**
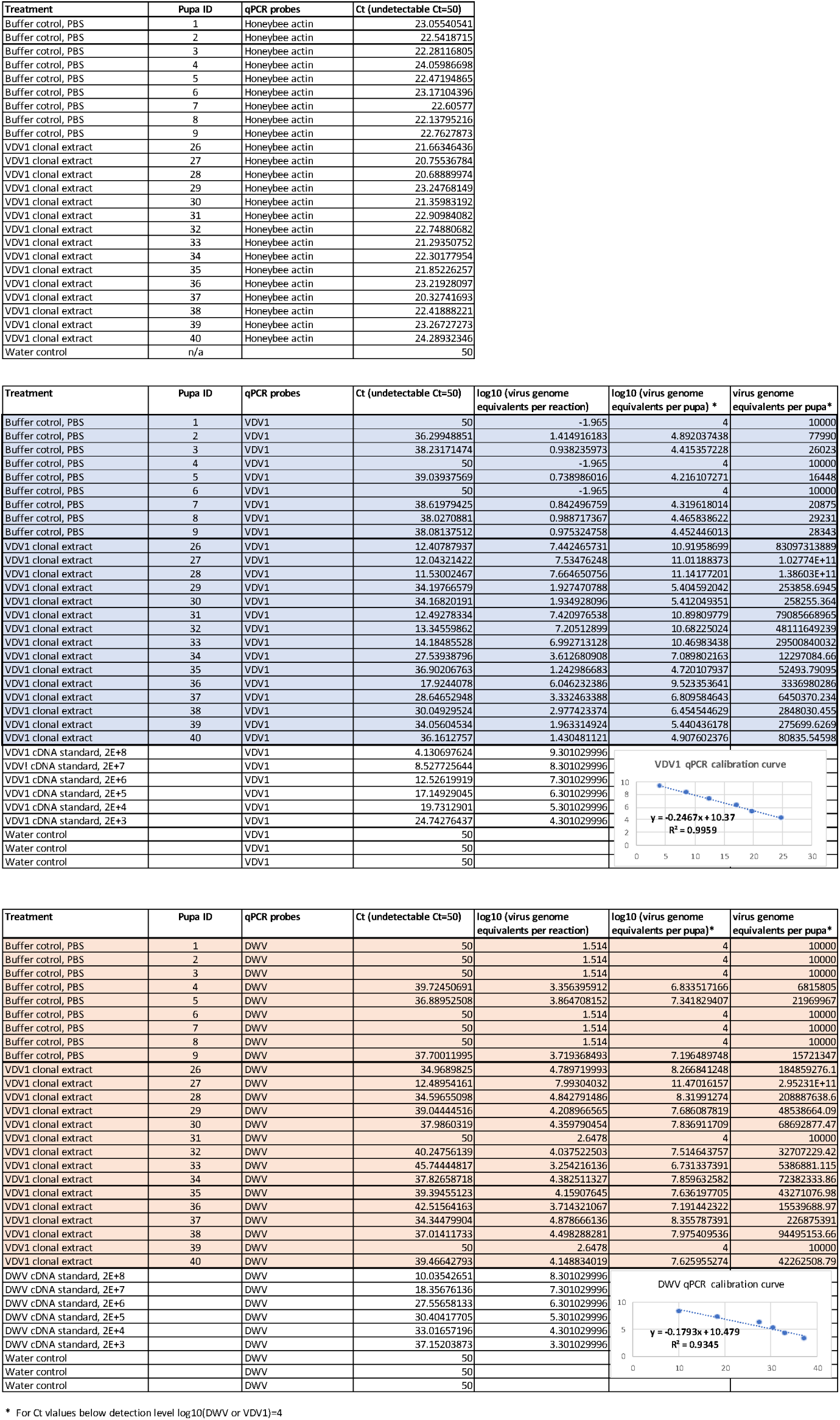
Virus acquisition and transmission experiment, RT-qPCR quantification of VDV1, DWV and honeybee actin.

**S1 Text. Design of the infectious cDNA clone of Varroa destructor virus-1 (VDV1) and production of the clone-derived VDV1 inoculum**

The full-length cDNA clone of Varroa destructor virus (VDV1) GenBank Accession number MN249174, was generated using total RNA from the honey bees sourced in California in 2016, isolate CA-07-2016, which showed high VDV1 and low DWV levels [1], using approach described in [2]. Two overlapping VDV1 cDNA fragments corresponding to the 5’ and 3’ sections of genomic RNA were amplified by RT-PCR using Superscript III reverse transcriptase (Invitrogen) and proof-reading Phusion DNA polymerase (New England Biolabs) to minimize amplification errors. The 5’ VDV1 cDNA fragment, positions 277-6333 nt, was generated using “Pr-VDV1-6300R” as a reverse transcription primer and “VDV1NotFseAsiSi-F” and “Pr-VDV1-6300R” as PCR primers (S1 Table). The 3’ VDV1 cDNA fragment, positions 4925-10148, was generated using “VDV1-AsciPmeI-A27R” as a reverse transcription primer and “Pr-VDV1-4800F” and “VDV1-AsciPmeI-A27R” as PCR primers, the primer sequences are given in Supplementary Table 1. These RT-PCR fragments containing overlapping 5’ and 3’ parts of VDV1 cDNA were cloned into the plasmid vector pTOPO-XL vector (Invitrogen) according to the manufacturer’s instructions to produce constructs pVDV1-12 and pVDV1-3, correspondingly. The cloned cDNA inserts were Sanger-sequenced to confirm integrity of the protein coding sequences and homology with the previously sequenced VDV1 isolates. The *Not*I-*Hind*III 4.9 kb fragment of pVDV1-12 was inserted into the *Not*I-*Hind*III-digested pVDV1-3 to produce a construct containing nearly full-length VDV1 cDNA clone, positions 277-10148 nt. Unique restriction sites *Not*I and *AsiS*I, which were introduced at the 5’ end of the cloned cDNA, were used to insert synthetic DNA sequence “T7-Rib-VDV1-5end” corresponding to the first 284 nucleotides of VDV1 preceded by the T7 RNA polymerase promoter and the ribozyme sequences [3] The resulting plasmid construct pVDV1-4, which contained introduced restriction site *AsiS*I in the IRES sequence at the position 277 nt, was linearized with *Pme*I restriction enzyme at the site located downstream the 3’ terminal polyA sequence to generate the template for *in vitro* transcription. The *in vitro* transcription was carried out with T7 RNA polymerase (HighScript, New England Biolabs) for 3 hours at +37°C, the template plasmid DNA was removed by digestion with RNAse-free Turbo DNAse (Ambion), the full-length VDV1 transcripts were purified by RNeasy column (Qiagen).

To produce clone-derived VDV1 inoculum, 5 μg of the purified *in vitro* RNA transcript in 10 μL of phosphate-buffered saline (PBS), was injected into the hemolymph of purple eye honeybee pupae using syringes with a 0.3 mm 31G hypodermal needle G31 (BD Micro-Fine).

The injected pupae were incubated 4 days at +33°C to allow development of the clone-derived virus infection and then were used to prepare tissue extracts containing the clone-derived VDV1 virus particles. The extracts were produced by homogenizing individual pupa with 1 mL of PBS, subjecting the homogenate to three cycles of freezing and thawing, clarifying extract by low-speed centrifugation (3000 rpm for 5 minutes), and filtering through a 0.22 μm nylon syringe filter. Concentrations of VDV1 and DWV was determined by qRT-PCR and identity of the clone-derived VDV1 was confirmed by sequencing and restriction digestion of the VDV1 IRES region.

---

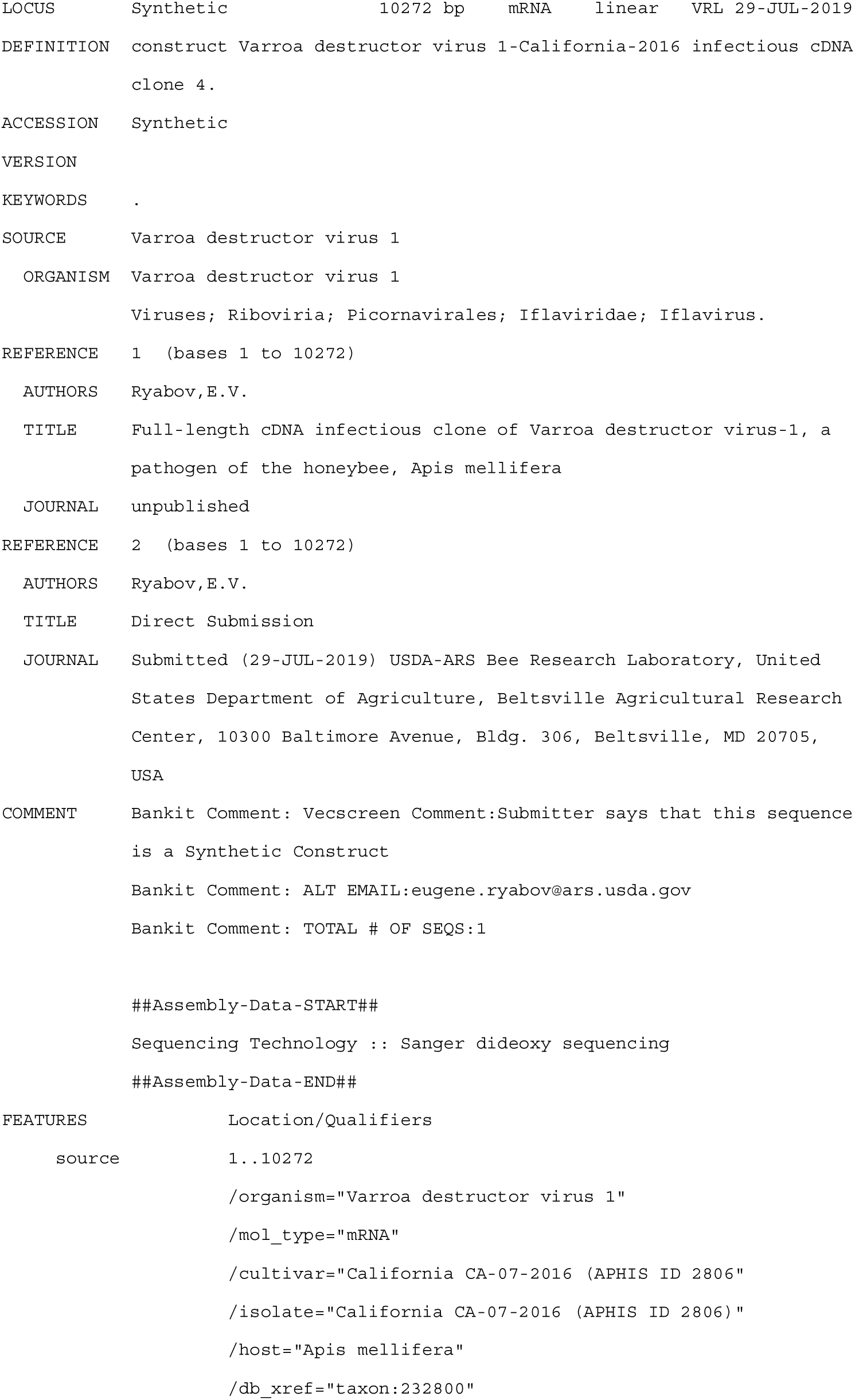
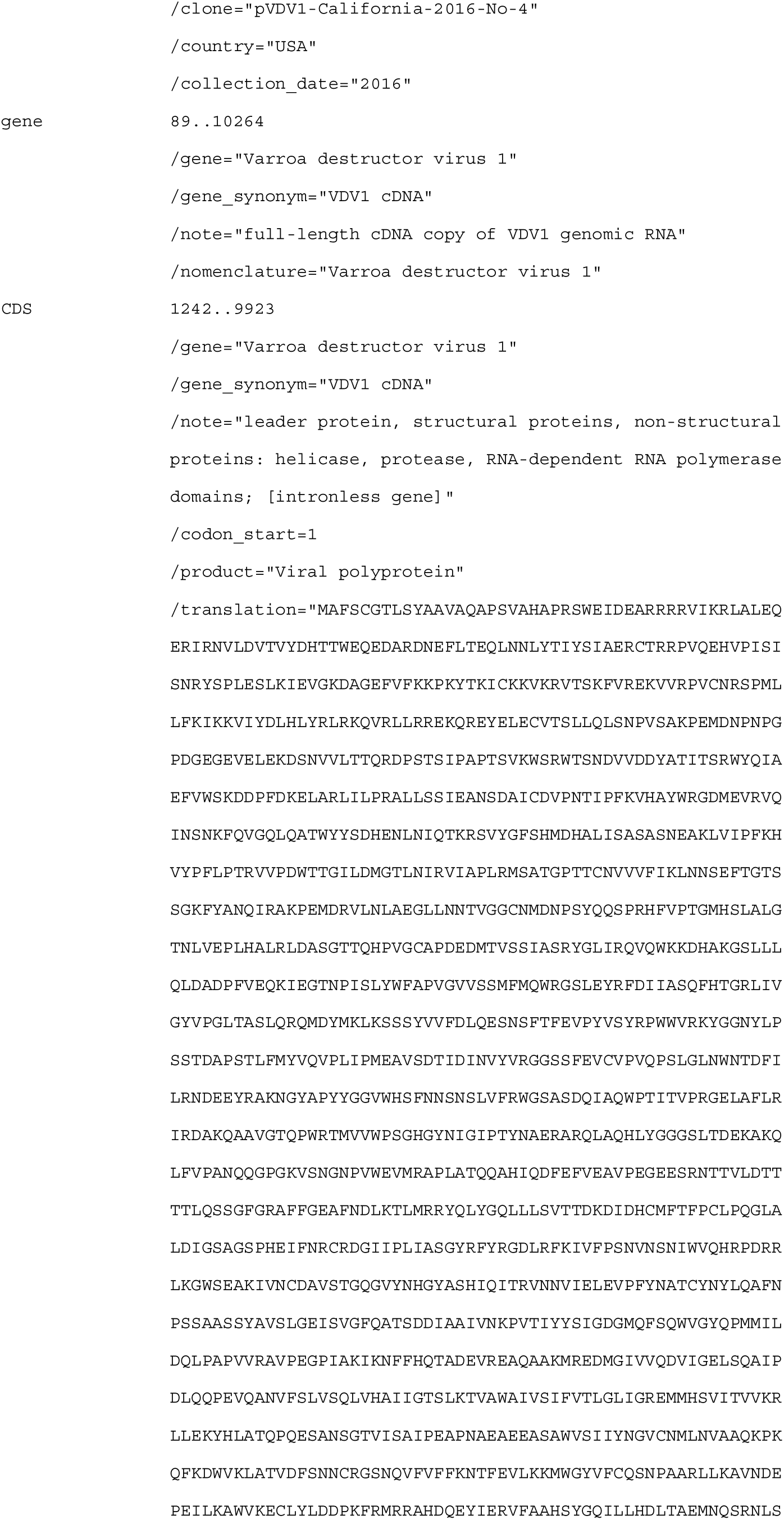
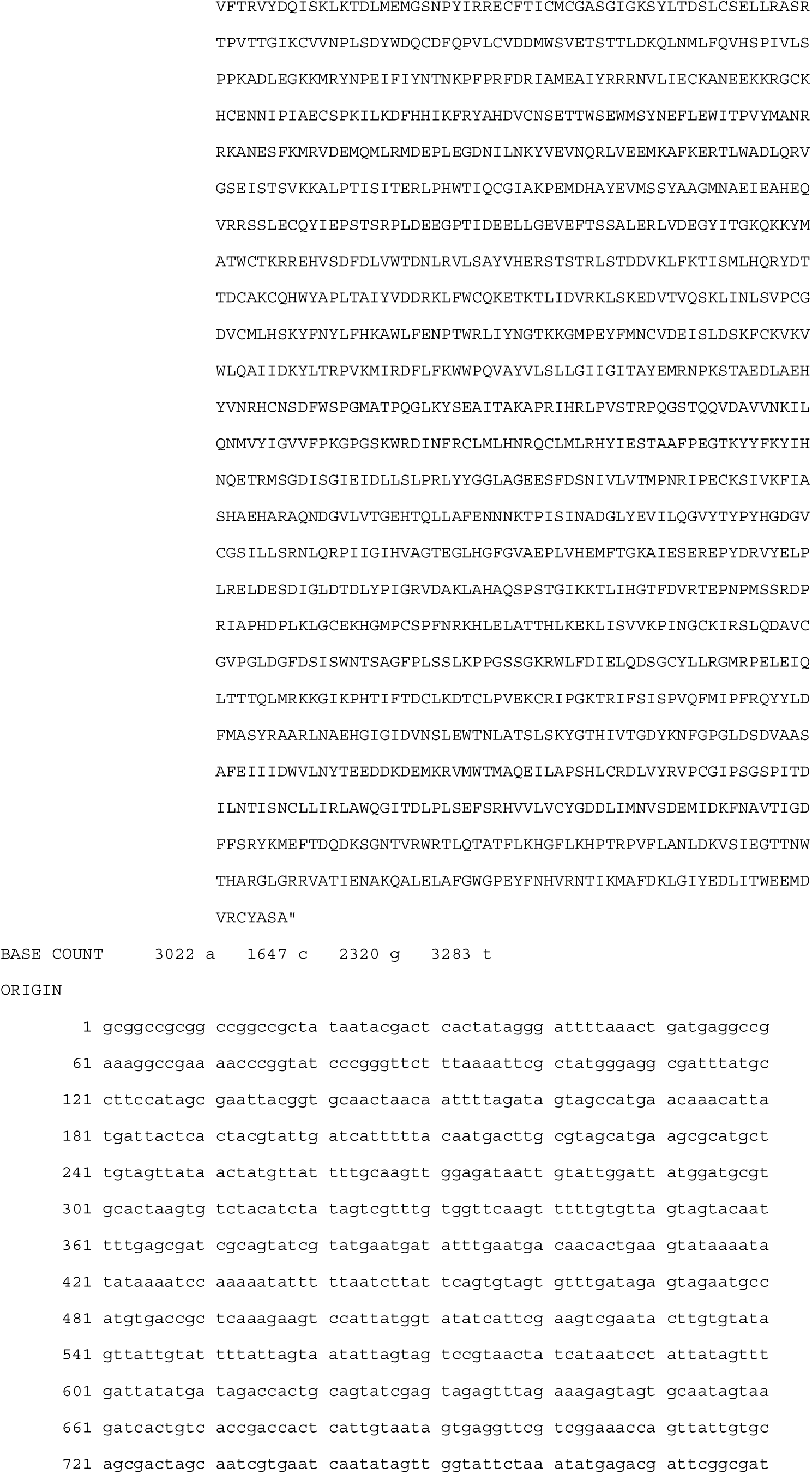
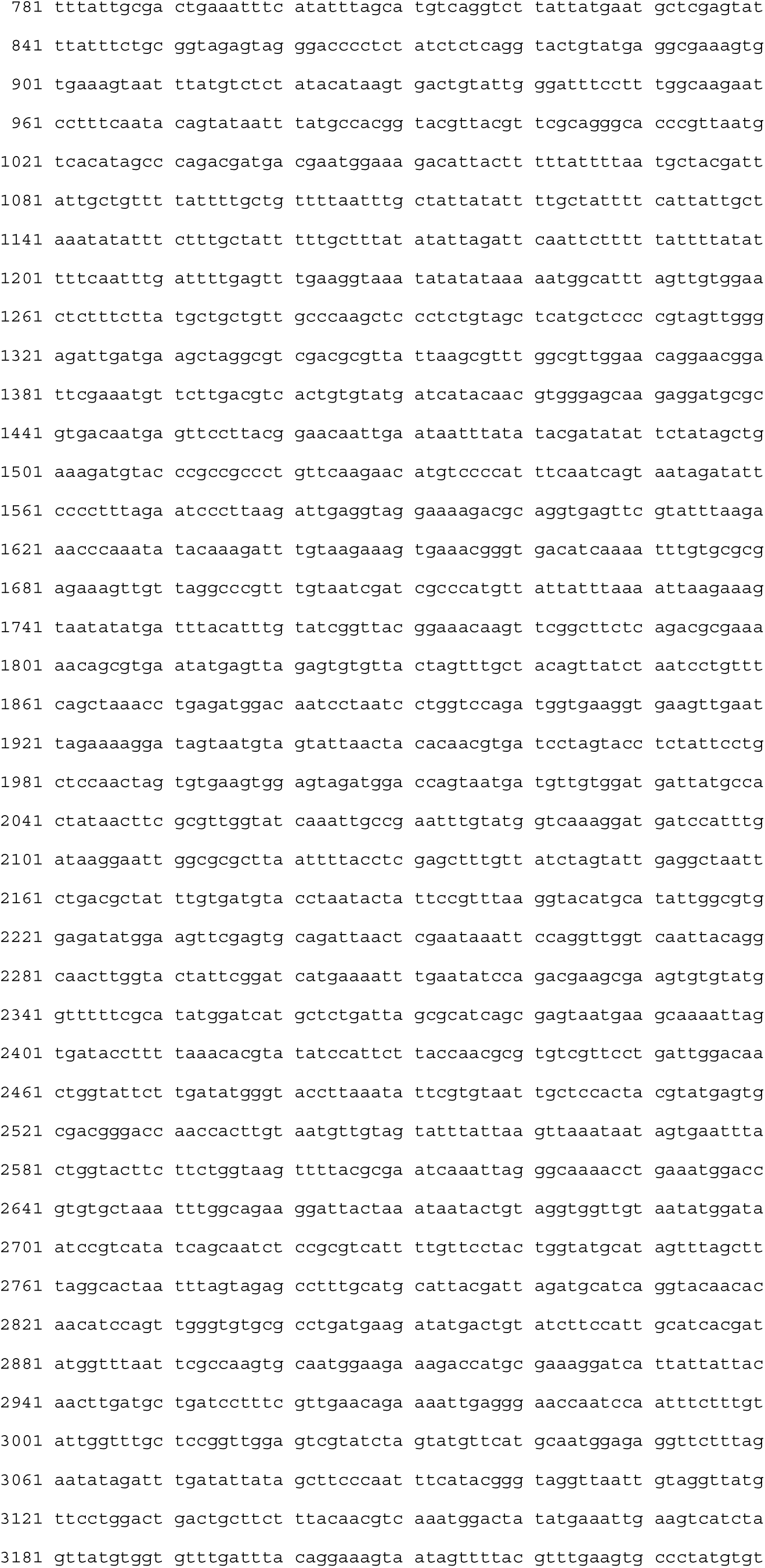
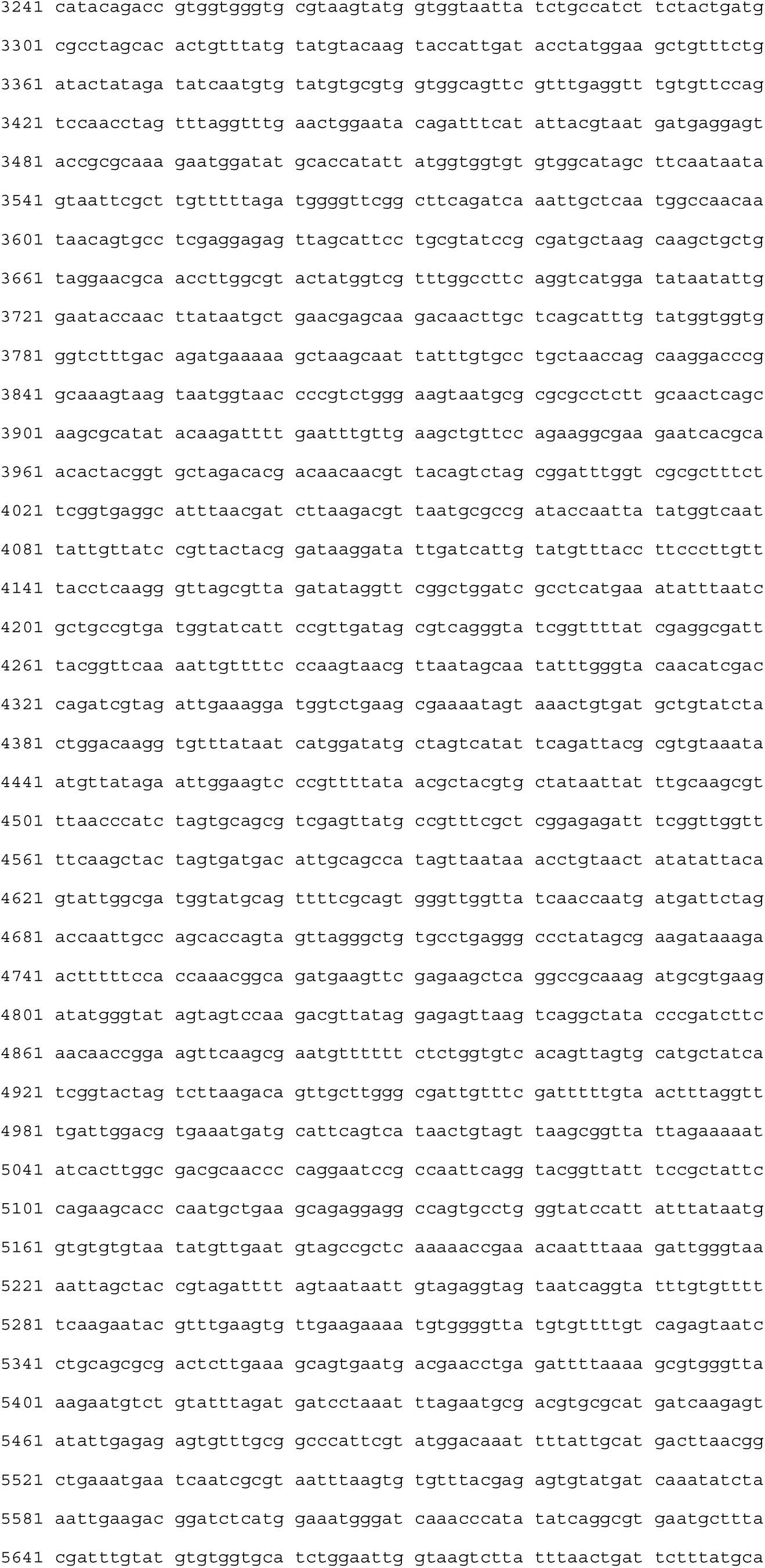
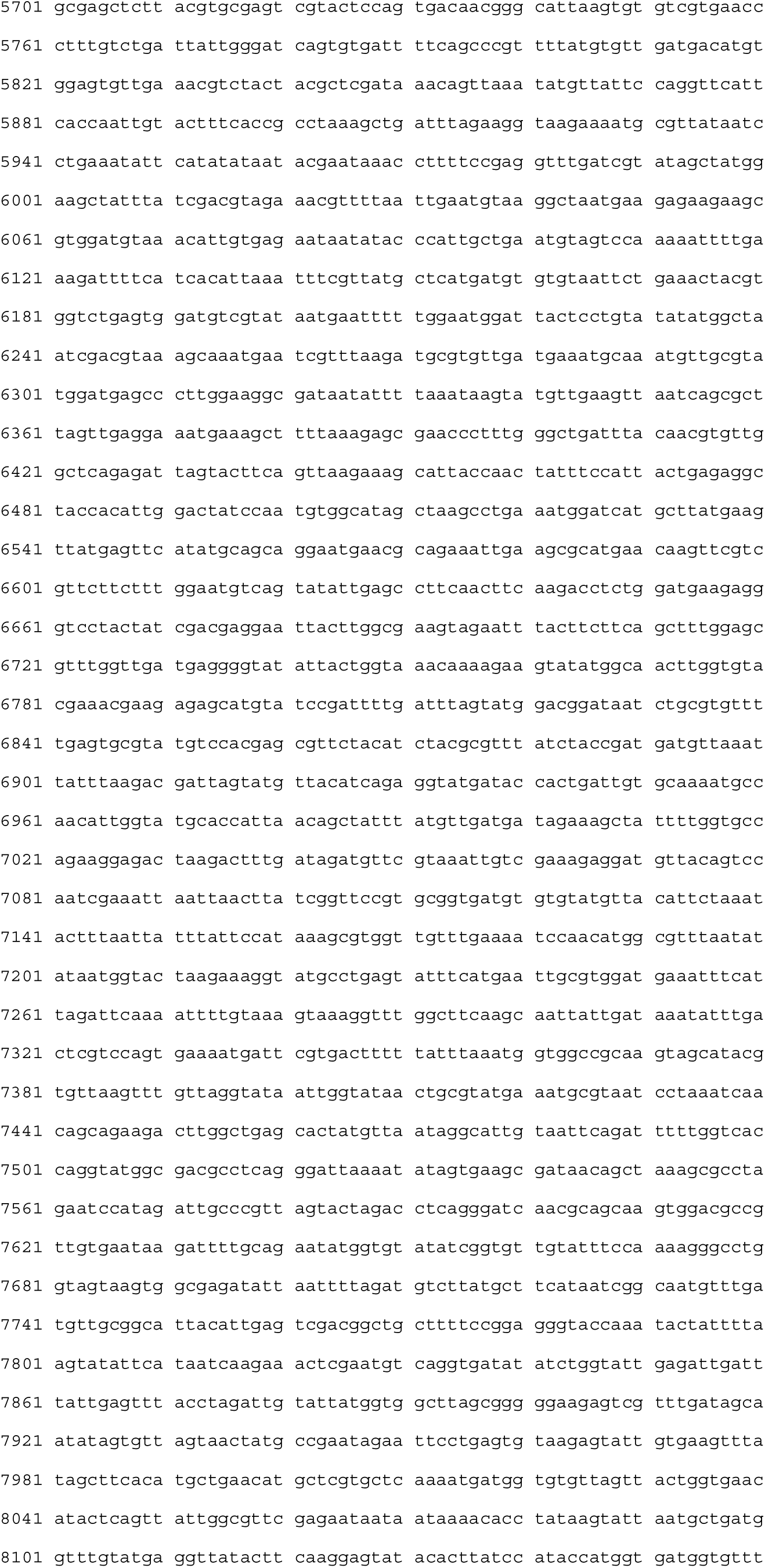
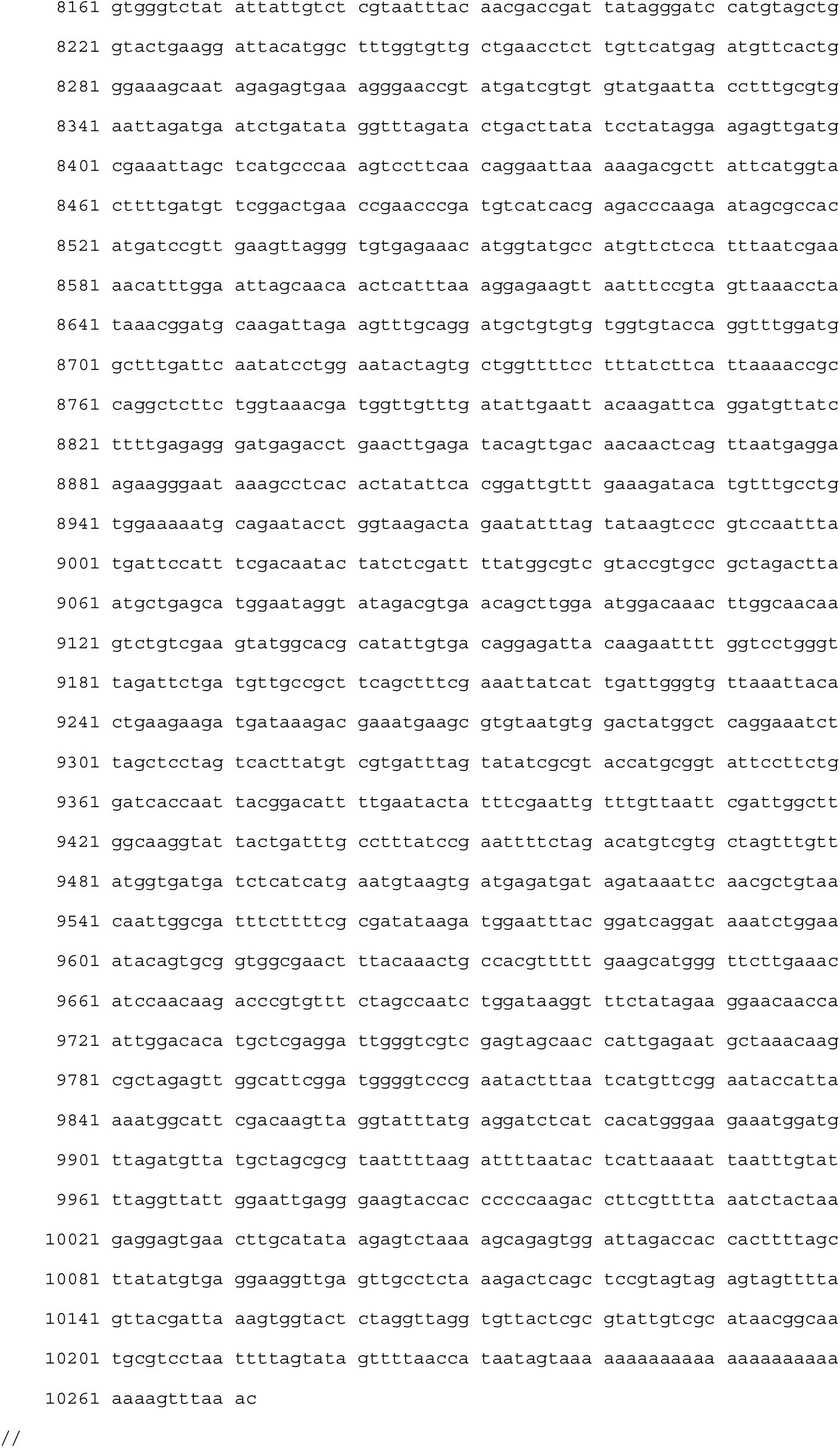

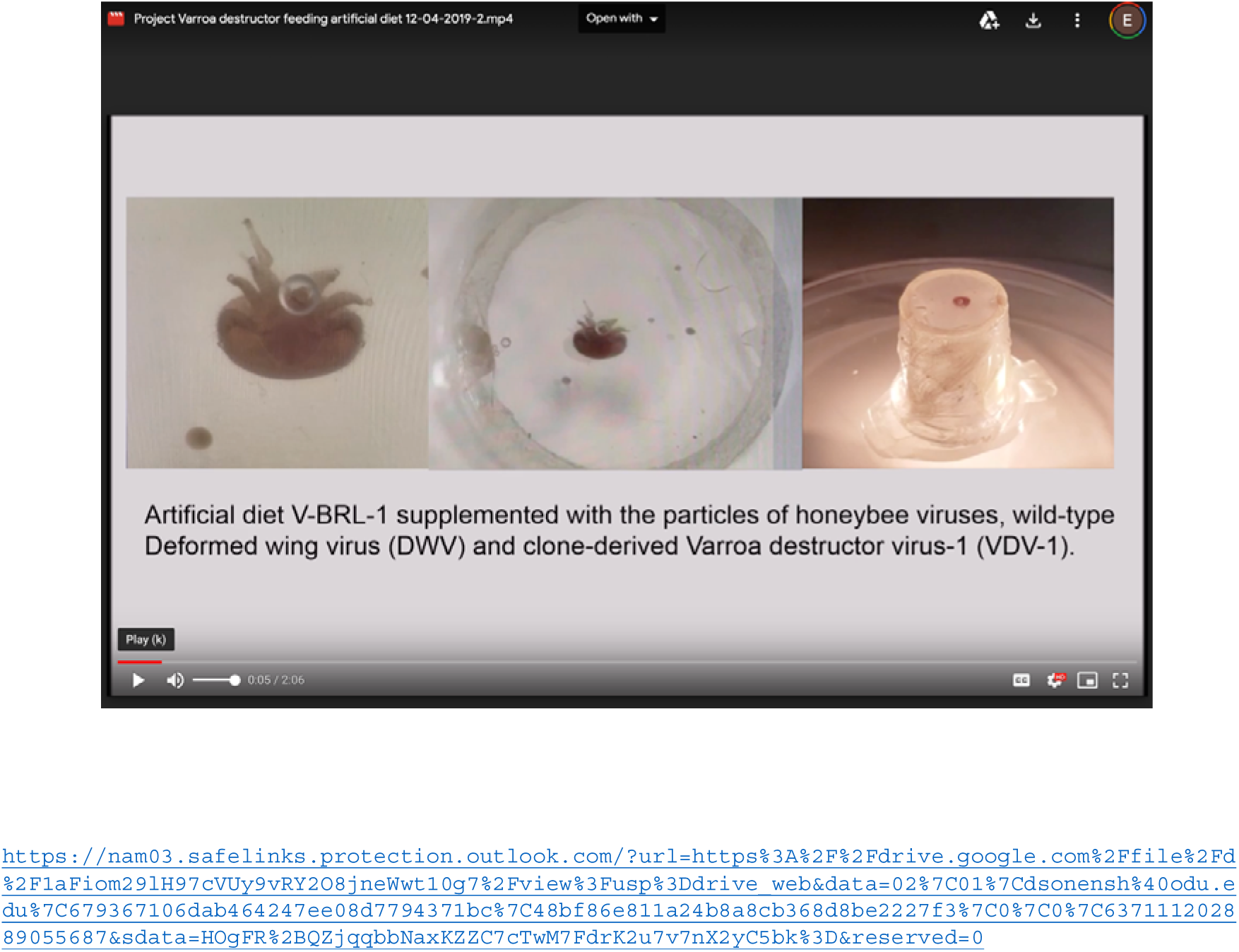

